# Holotomographic microscopy reveals label-free quantitative dynamics of endothelial cells during endothelialization

**DOI:** 10.1101/2024.11.04.621934

**Authors:** William D. Leineweber, Gabriela Acevedo Munares, Christian Leycam, Juliette Noyer, Patrick Jurney

## Abstract

Holotomograhic microscopy (HTM) has emerged as a non-invasive imaging technique that offers high-resolution, quantitative 3D imaging of biological samples. This study explores the application of HTM in examining endothelial cells (ECs). HTM overcomes the limitations of traditional microscopy methods in capturing the real-time dynamics of ECs by leveraging the refractive index (RI) to map 3D distributions label-free. This work demonstrates the utility of HTM in visualizing key cellular processes during endothelialization, wherein ECs anchor, adhere, migrate, and proliferate. Leveraging the high resolution and quantitative power of HTM, we show that lipid droplets and mitochondria are readily visualized, enabling more comprehensive studies on their respective roles during endothelialization. The study highlights how HTM can uncover novel insights into EC behavior, offering potential applications in medical diagnostics and research, particularly in developing treatments for cardiovascular diseases. This advanced imaging technique not only enhances our understanding of EC biology but also presents a significant step forward in the study of cardiovascular diseases, providing a robust platform for future research and therapeutic development.

## Introduction

Holotomographic microscopy (HTM), also known as quantitative phase tomography, is an emerging optical microscopy approach that enables label-free, quantitative imaging at high resolution and in 3D.^1–3^ It leverages optical interferometry to measure the phase shift of a coherent light wavefront passing through the sample, a parameter directly related to its refractive index (RI) and thickness distributions. By capturing multiple holographic images at varying illumination angles or sample orientations, HTM obtains projections analogous to those used in computed tomography, ultimately providing a quantitative, label-free visualization of subcellular architecture within living cells without the need for exogenous labeling steps. Much of the groundwork has been laid in recent years to associate cellular components to their RI, such as lipid droplets and mitochondria.^4–7^ Biological insights arising due to these unique measurements include tracking the transfer of cholesterol from host to pathogen ^8^, observing multi-organelle rotation patterns within HeLa cells ^7^, and differentiating between cell types in co-cultures.^9^ These applications demonstrate that HTM is well-suited to study cellular processes involving dynamic behaviors that are otherwise challenging to measure with conventional imaging approaches.

Endothelialization - the process by which endothelial tissue is formed - is an active area of research in cardiovascular disease. Injured endothelium contributes to disease progression, and surgical interventions often require implantation of devices which require some portion to be endothelialized. Poor endothelialization is a leading cause of implant failures, leading to worse patient outcomes and increases in costs ^10–12^. *In vitro* assessments of endothelialization often focus on how different biomaterials or growth factors impact endothelial cell (EC) attachment, spreading, migration, or proliferation.^13,14^ The ECs used in these assays are often primary cells, making them challenging to culture, engineer, or label. Due to these limitations, the readouts of EC function often rely on end-point analyses that may miss live-cell dynamics relevant to endothelialization progression.

Understanding the nuances of endothelialization is vital for improving cardiovascular therapies, particularly in the context of vascular grafts, stents, and other medical devices. In this study, we used HTM to measure EC dynamics during the early stages of endothelialization. We hypothesized that the unique capabilities of HTM to provide high-resolution, three-dimensional, label-free images would generate novel insights into EC behavior during endothelialization. We found that HTM could be used to observe ECs for long imaging windows without introducing artifacts from dyes or phototoxicity associated with fluorescence microscopy. The quantitative nature of HTM, particularly the measurement of RI values, added valuable dimensions to cellular analysis that correlate with cellular states and processes, such as attachment and migration. We further showed that combining HTM with immunofluorescence imaging augmented the information gained from either modality separately. Overall, we found that HTM is particularly advantageous for studying endothelialization, where the interplay of cellular morphology, behavior, and interactions with the substrate are critical.

## Results

### Holotomography Reveals Multiscale Structures in Endothelial Cells

The instrumentation (Fig. 1A) typically includes a low-coherence light source, beam-splitting optics, and a digital holographic camera system configured so that a reference beam interferes with the beam transmitted through the sample. As the illumination angle or sample orientation is systematically varied, multiple interferograms are recorded and processed using computational algorithms rooted in diffraction tomography and inverse scattering theory. The resulting three-dimensional refractive index map enables differentiation of organelles and cellular structures based on their inherent optical properties (Fig. 1B). Because the refractive index correlates with biomolecular density, HTM reveals subtle morphological details and dynamic cellular processes, making it ideally suited for studying endothelialization and other complex biological phenomena.

**Figure 1:**
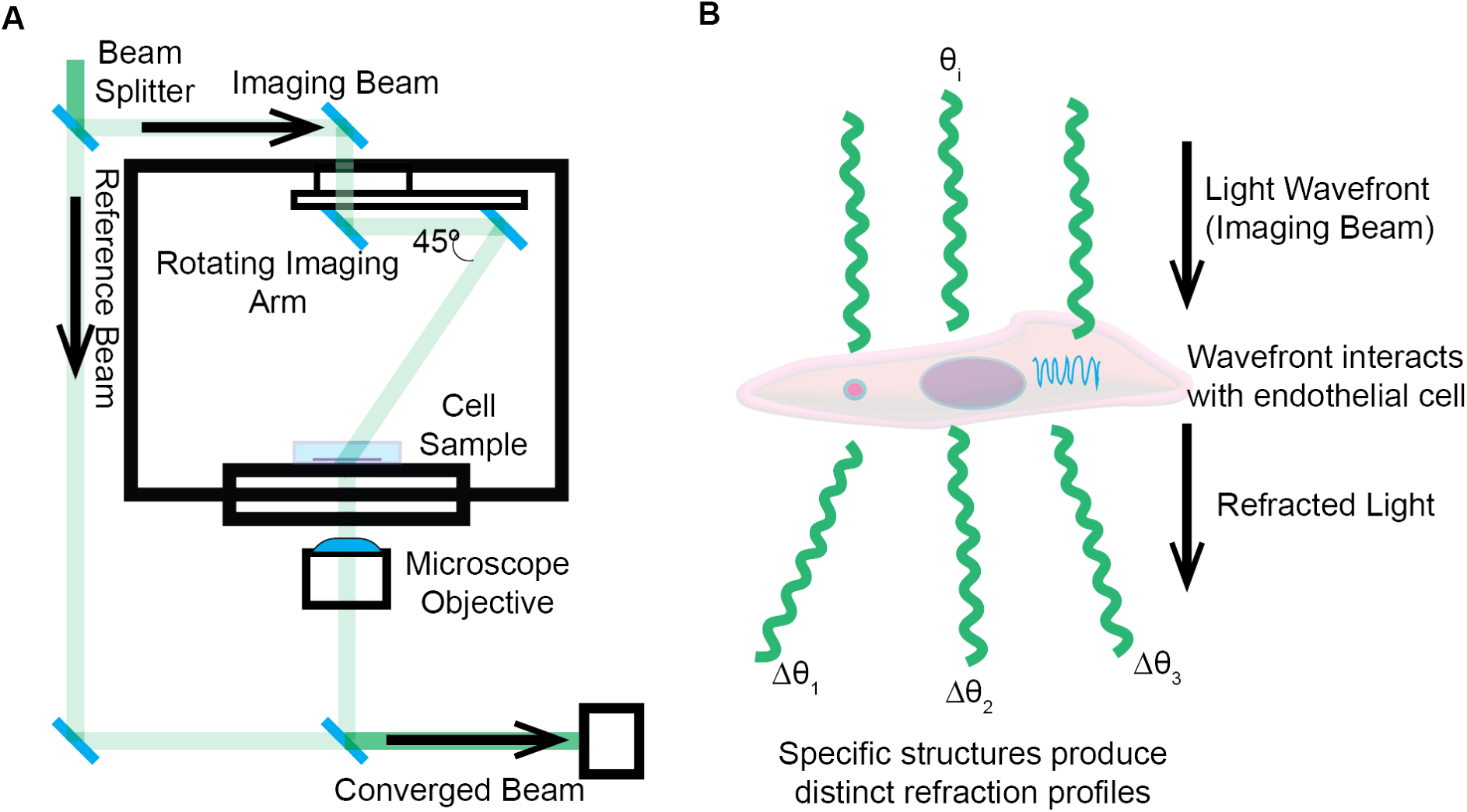
Principle of holotomographic microscopy for label-free, quantitative imaging. **(A)** Schematic of a typical HTM setup. A low-coherence light source is split into an imaging beam that traverses the sample and a reference beam. As the angle of illumination or sample orientation changes, multiple interferograms are recorded. These are computationally reconstructed into a three-dimensional refractive index (RI) map, providing quantitative, label-free contrast. **(B)** Conceptual illustration of an endothelial cell demonstrating how subcellular compartments, such as organelles, differ in their RI. These RI differences enable HTM to visualize cellular structures without external dyes, capturing subtle morphological details relevant to processes like endothelialization.

We sought to determine the utility of HTM in imaging ECs during the early stages of endothelialization. Our primary interest was to assess whether this approach could provide high-resolution, label-free images that reveal multiple cellular compartments across multiple size scales. Visualization of ECs as 3D reconstructions (Supplemental Video S1) provided a useful topographical view of the main cell bodies, while max intensity projections revealed rich subcellular detail (Fig. 2 A). When imaging a field of view with multiple cells, HTM imaging clearly distinguished individual cell boundaries and their nuclei (Fig. 2 B). By zooming into a field of view with four to five ECs, cellular protrusions such as lamellipodia and filopodia were readily observable (Fig. 2 C). Cell-cell interactions were also captured, including zipper-like cell junctions and tunneling nanotubes (Fig. 2 D). At this cellular resolution, the spatial distribution of organelles was clear enough to determine colocalization patterns. Zooming in further, highly-resolved subcellular compartments could be visualized, including individual lipid droplets, nucleoli, extracellular vesicles, and mitochondria (Fig. 2 E). These findings demonstrate that HTM can simultaneously visualize multiple subcellular compartments in high resolution and without labeling, providing detailed insights into cellular structures involved in endothelialization.

**Figure 2:**
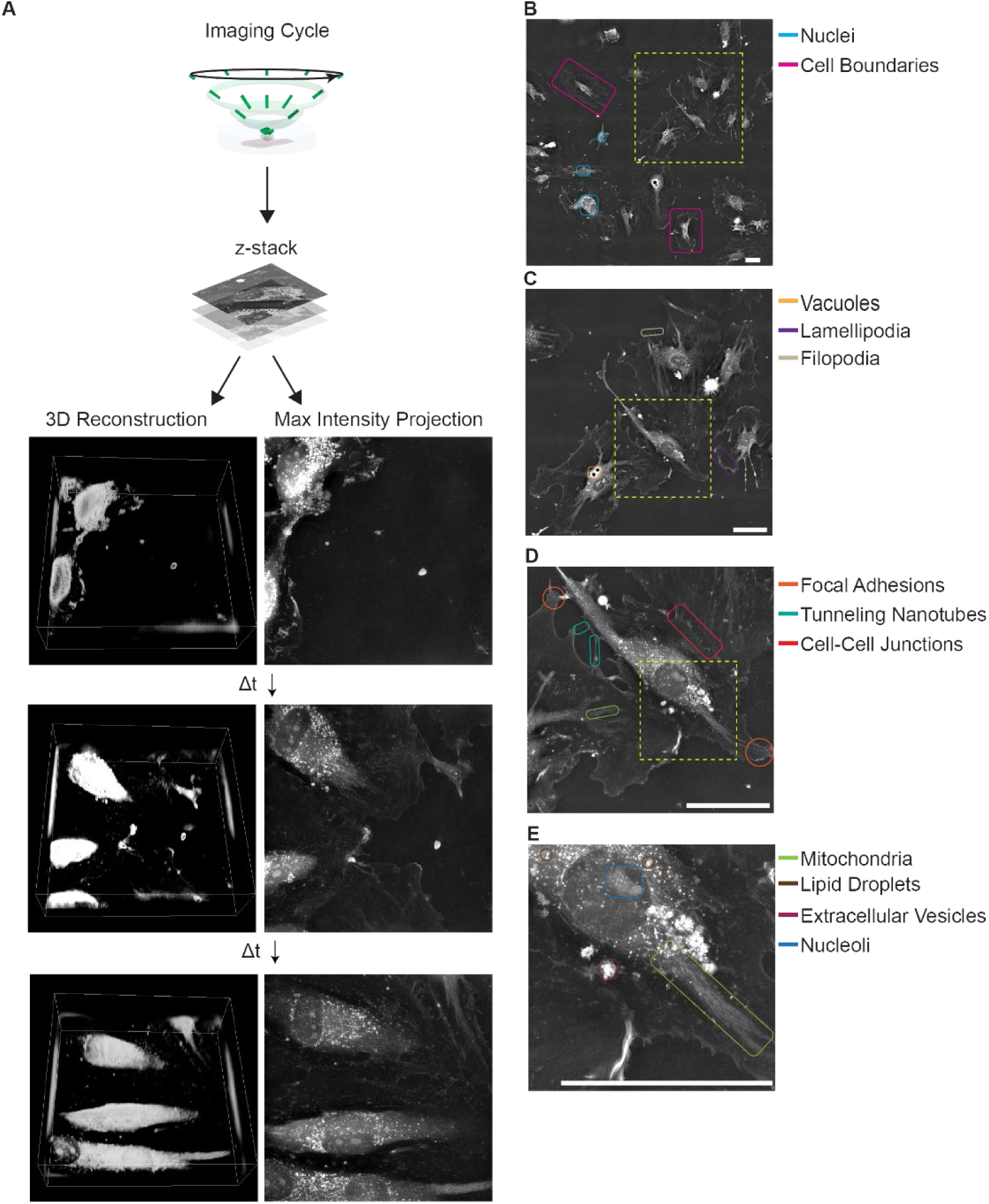
Multiscale visualization of endothelial cell architecture and behavior using holotomographic microscopy. **(A)** Representative holotomographic images of HUVECs at 5 minutes, 10 hours, and 20 hours post-seeding, showing their progression during endothelialization. The HTM imaging cycle involves acquiring a 3D RI map from multiple illumination angles, which can be rendered as a three-dimensional reconstruction or maximum intensity projection (MIP). **(B)** Large-field tiled acquisitions highlight nuclei and clear cell boundaries, enabling the simultaneous observation of numerous cells. **(C)** A higher-magnification view of a small neighborhood reveals vacuoles, lamellipodia, and filopodia, providing detailed subcellular context. **(D)** Further zooming in detects focal adhesions, tunneling nanotubes, and cell-cell junctions, illustrating the complexity of intercellular interactions. **(E)** At the highest resolution, mitochondria, lipid droplets, extracellular vesicles, and nucleoli are visualized label-free, demonstrating the ability of HTM to identify multiple organelles and subcellular compartments. Scale bar = 20 μm.

### Combining Holotomography with Fluorescence Imaging Enhances Cellular Component Identification

The cellular features captured by HTM imaging provide generalizable, label-free readouts of cellular components but lack specificity at the protein composition level. To overcome this limitation, we combined immunofluorescence with HT, enabling direct comparisons of protein localization with the label-free features observed in HT images. To illustrate this approach, we examined cell-cell contacts during endothelialization by staining for vascular endothelial cadherin (VE-cad), a key marker for adherens junctions that develop during this process. VE-cad expression increased over time, and localized to EC-EC contacts robustly within four hours of seeding HUVECs (Fig. 3 A). Overlaying VE-cad fluorescence images with the corresponding HT micrographs allowed us to assess the relationship between RI values at cell borders and VE-cad levels. While RI values often highlighted cell borders due to differences in RI between adjacent cells, quantitative analysis revealed no significant correlation between VE-cad localization and RI values at these borders at the 12-hour mark (Pearson’s r-value = −0.02, Costes P-value = 0.36) (Fig. 3 B). These findings highlight the orthogonality of HTM and immunofluorescence in visualizing cell-cell interactions.

**Figure 3:**
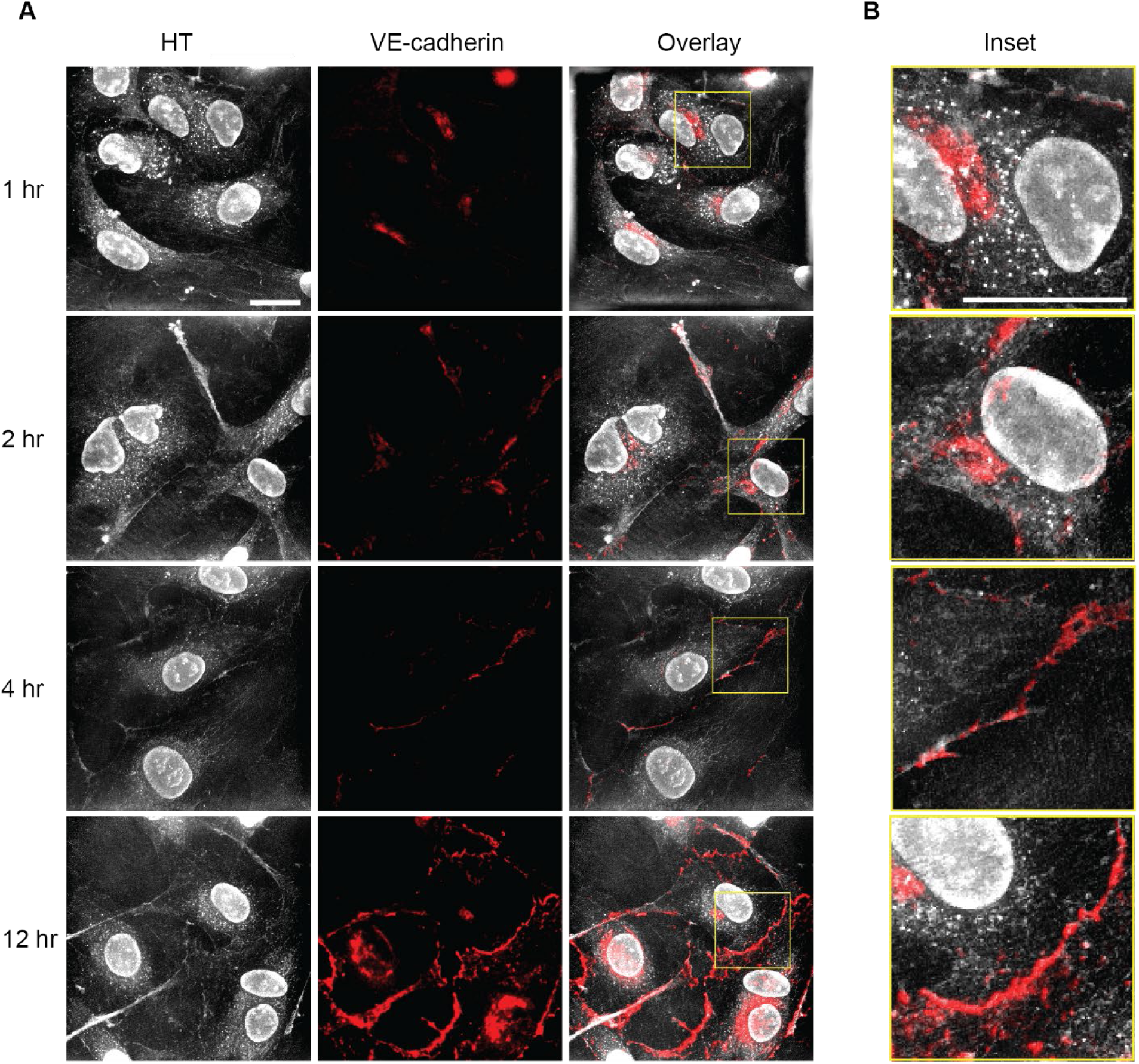
Combining immunofluorescence with holotomographic microscopy provides complementary insights into endothelial cell structure. **(A)** Representative holotomographic (HT) and immunofluorescence (IF) images of HUVECs stained for VE-cadherin at various timepoints post-seeding. The fluorescence signal highlights the progressive increase and localization of VE-cadherin along cell-cell boundaries, while the HT images provide label-free structural context. **(B)** Higher magnification views of merged HT and IF channels show that VE-cadherin distribution does not strongly correlate with refractive index (RI) variations. Despite the presence of distinct morphological features in the HT images, VE-cadherin localization is functionally independent of these RI-based contrasts. Scale bar = 20 μm.

### Live-Cell HTM Time-Lapse Imaging Reveals Dynamic Refractive Index Changes During Endothelialization

HTM is compatible with live-cell time-lapse imaging, and its quantitative nature allows RI values to serve as indicators of cell states. We imaged HUVECs adhering and migrating shortly after seeding and quantified their mean RI values over time (Supplemental Video S2). The ECs displayed dynamic morphology and movement over the 16-hour acquisition, indicative of the early stages of endothelialization (Fig. 4 A). Cells I and II were already attached to the substrate at the beginning of the acquisition, while cells III - VI adhered and spread out during the imaging period. The average RI values of the cells were measured for each frame of the time lapse following manual segmentation of cell boundaries (Fig. 4 B, Supplemental Video S3).

**Figure 4:**
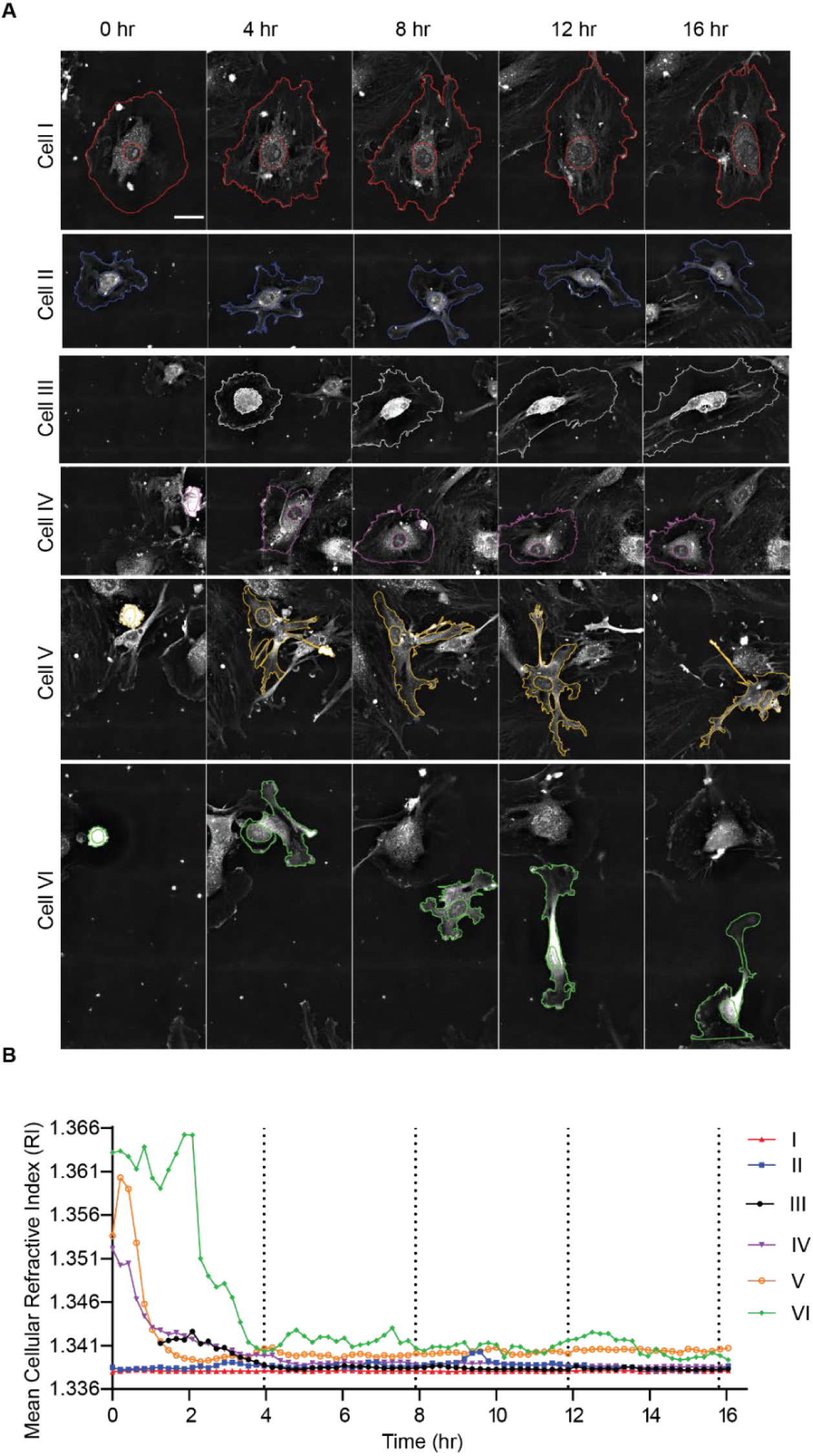
Live-cell holotomographic time-lapse imaging captures endothelial cell heterogeneity. **(A)** Representative HT images of individual HUVECs monitored over a 16-hour period, with outlines of nuclei and cell boundaries overlaid to emphasize dynamic morphological changes as cells adhere, spread, and migrate. Scale bar = 20 μm. **(B)** Quantification of the mean refractive index (RI) for each cell throughout the time-lapse reveals distinct, cell-specific RI trajectories, underscoring the heterogeneity of early endothelialization processes.

We observed three notable trends in the RI values corresponding with relevant endothelialization processes: 1) Cell anchoring, attachment, and spreading coincided with a dramatic decrease in RI values. 2) Cells appeared to reach an equilibrium RI value after sufficient time. 3) Motile cells maintained a higher mean RI value even after the initial cell attachment and anchoring phases. These results demonstrate that RI measurements obtained via HTM can reflect dynamic changes in ECs during endothelialization. We next sought to further explore how the cell RI values related to morphological and functional changes to ECs.

### Quantitative Analysis of Cell Morphology and RI Values During Early Endothelialization (*Fig. 5*)

The initial steps of endothelialization involve ECs capturing, tethering, activating, arresting, and adhering to a substrate (Fig. 5 A, B). A prominent observation of HTM imaging was a precipitous drop in cellular RI values, followed by stabilization during anchoring, attachment, and spreading (Fig. 4 B). Utilizing the high-resolution images, we obtained accurate measurements of cell area (Fig. 5 C). Changes in cell area served as useful indicators of cell activation and arrest; however, RI values provided a more consistent readout, being less influenced by variability in cell shape that can introduce noise into area measurements. RI values were highly sensitive to changes in cell area for small cells, rapidly decreasing as cell areas increased (Fig. 5 D). As the ECs spread out more and stabilized into monolayers, their RI values also stabilized, providing an insight into the maturity and stability of the endothelial monolayer being imaged. This relationship meant that time until cell anchoring could be easily readout by measuring the RI curve prior to stabilization. Some cells anchored within the first hour, while others did not attach until closer to four hours.

**Figure 5:**
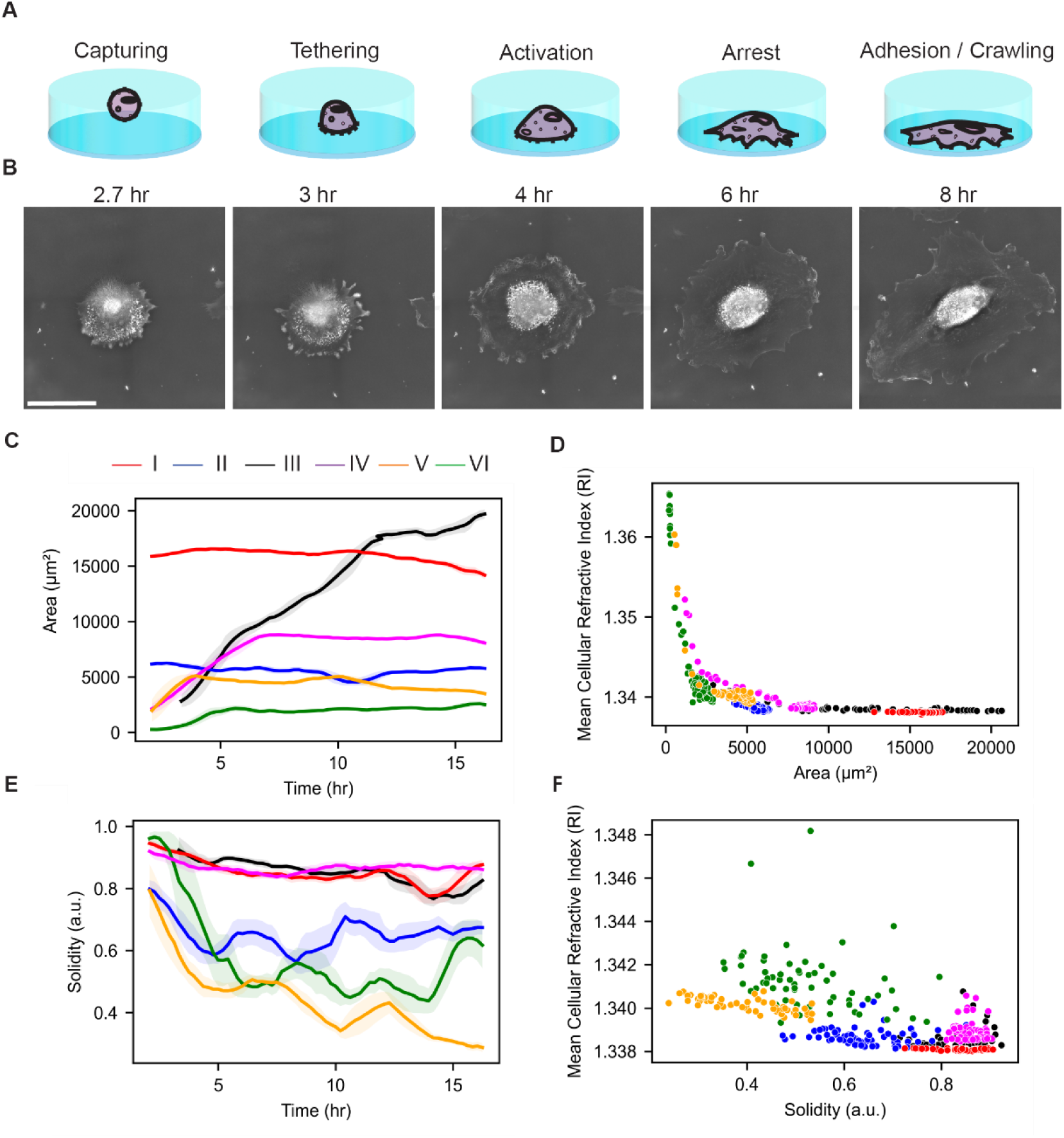
Quantitative characterization of endothelial cell anchoring and attachment using holotomography. **(A)** Schematic illustrating early endothelialization stages: cells are initially captured by weak interactions, then form stronger adhesions during tethering, become activated, arrest, and eventually achieve stable attachment before migrating. **(B)** Representative HT images of a single HUVEC undergoing these progressive stages, highlighting morphological changes over time. **(C)** Time-lapse measurements of cell area for individual HUVECs show increasing spread as cells progress through early endothelialization. **(D)** Mean refractive index (RI) decreases with increasing cell area, particularly when cell areas are below 100,000 µm², reflecting changes in cellular density during spreading. Each point represents a single cell at a given time point. **(E)** Cell solidity values generally decrease or remain stable in the initial hours, indicating shifts in cell shape as they attach. Lines represent mean values smoothed over a two-hour window; shaded regions indicate the 95% confidence interval. **(F)** After the initial capture and tethering phase (t > 4 hr), cells with lower solidity values exhibit higher RI, revealing a negative correlation between cell shape complexity and optical density. Each point corresponds to a single cell at a given time point after the initial phases.

After establishing the correlation between cell area and RI values (Spearman r = −0.717, p = 1.62e-13), we investigated how cell shape is associated with endothelialization. We quantified cell shape using solidity, where values closer to one indicate convex cells and values closer to zero indicate more irregularly shaped cells. During the first four hours after seeding, we observed sharp decreases in solidity for the ECs during capturing and tethering phases (Fig. 5 E). Notably, whereas the areas of cells V and VI stabilized within the first four hours, their solidity values remained variable and gradually decreased over time (Fig. 5 E). For timepoints after cells had adhered (after the initial fo7ur hours), we observed that cells with lower solidity values typically exhibited higher RI values (Spearman r = −0.652, p = 4.79e-48 (Fig. 5 F). These results demonstrate that RI measurements correlate with changes in cell area and solidity during early endothelialization.

### Correlation Between Cell Motility and Refractive Index in Endothelial Cells

An additional advantage of high-resolution, label-free time-lapse HTM is the ability to perform cell migration analyses during endothelialization. As observed earlier, ECs with lower solidity values tend to have higher RI values (Fig. 5 F). We hypothesized that these cells are also more motile, given that their shapes are typically associated with more migratory cells.

EC migration can be broken down into the stages of sensing, extension, attachment, contraction, and rear release (Fig. 6 A). Using HTM, we tracked the movement of individual cells by monitoring their center of mass without the need for additional dyes or labeling techniques. The clear cell outlines provided by HTM facilitated accurate tracking. The trajectories of individual cells exhibited heterogeneous behaviors; some cells displayed highly persistent migration, while others appeared more random (Fig. 6 B). Quantitative analyses using mean squared displacement (MSD) measurements clearly showed three motile cells (Cells IV, V, and VI) and three non-motile cells (Cells I, II, and III) (Fig. 6 C). Instantaneous cell speeds showed cyclical increases and decreases, with varying periods and amplitudes unique to each cell (Fig. 6 D). Notably, Cells V and VI maintained higher RI values over the entire time course, even after attachment to the substrate. Cell IV also migrated a similar net distance as these two cells, though much of this migration occurred at earlier time points while the RI values were higher. A positive correlation between cell speed and RI value was observed (Spearman r = 0.28, p = 2.36e-8) (Fig. 6 E). These findings indicate that higher RI values are associated with increased cell motility during endothelialization.

**Figure 6:**
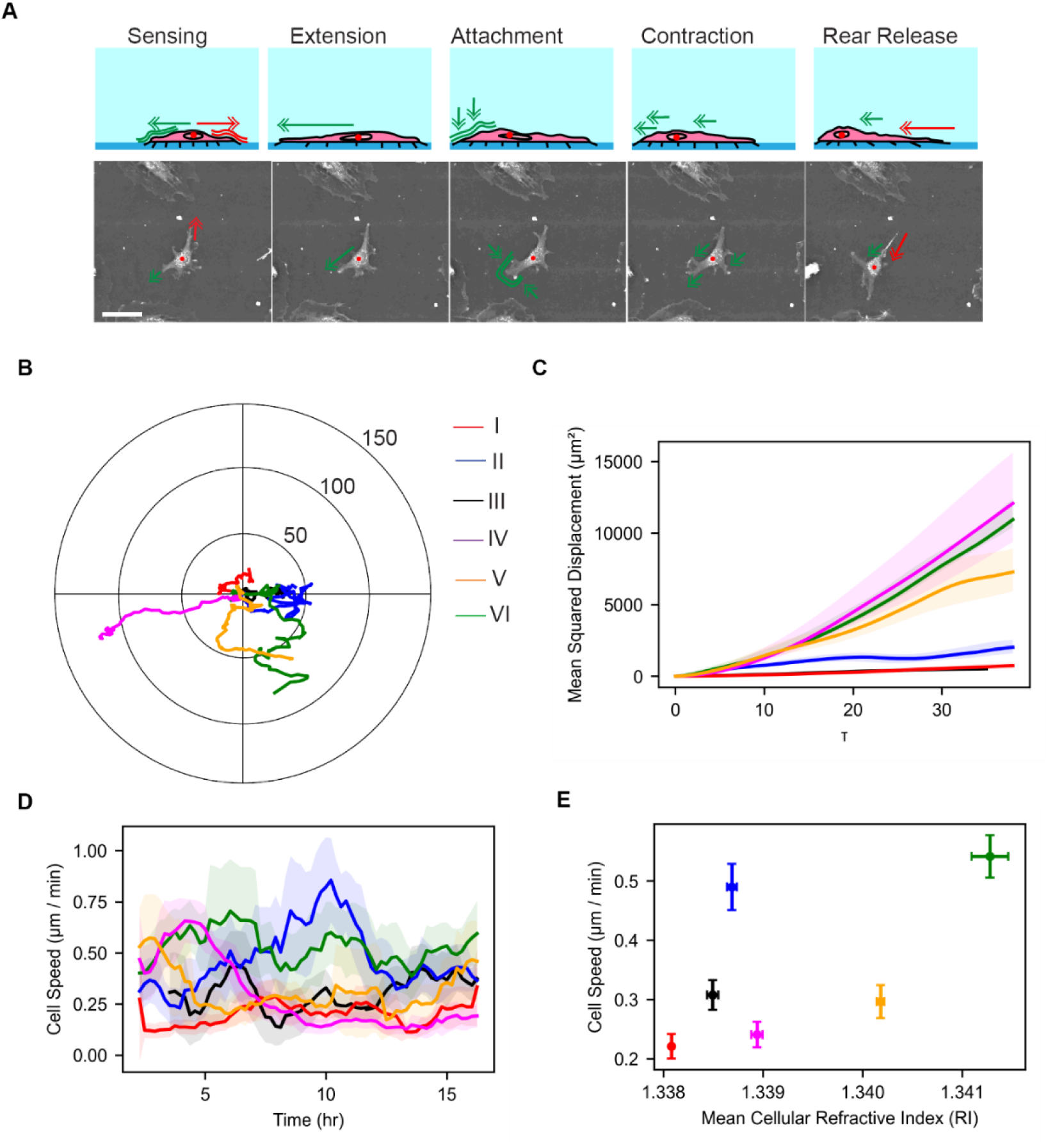
Quantifying endothelial cell migration and its relationship to refractive index using holotomography. **(A)** Schematic (top) illustrating the cyclical stages of endothelial cell migration—environmental sensing, protrusion formation, adhesion, contraction, and rear release—followed by representative HT images (bottom) of a single HUVEC undergoing these steps. Scale bar = 70 μm. **(B)** Individual cell trajectories tracked over 16 hours show heterogeneous migration behaviors within the population. **(C)** Mean squared displacement (MSD) analyses identify distinct subpopulations of migratory and non-migratory cells, with lines representing mean MSD values and shaded regions indicating standard error of the mean. **(D)** Instantaneous cell speed, averaged over two-hour intervals, fluctuates periodically and uniquely for each cell, highlighting dynamic, cell-specific migratory patterns. Shaded regions show the 95% confidence interval. **(E)** After the initial anchoring phase (t > 4 hr), a positive correlation emerges between mean cellular RI and cell speed, suggesting that higher RI values may be indicative of more motile endothelial cells. Each point represents a single cell’s mean RI and speed, with error bars reflecting standard error of the mean.

## Discussion

### Summary of Main Findings

In this study, we demonstrated the utility of HTM in visualizing ECs during the early stages of endothelialization. HTM provided high-resolution, label-free images across multiple scales, from whole-cell morphology to subcellular structures such as lipid droplets and mitochondria. By combining HTM with fluorescence imaging, we enhanced cellular component identification, showing that these techniques offer complementary, orthogonal approaches to visualizing cell-cell interactions. Our quantitative analyses revealed RI measurements obtained via HTM reflect dynamic changes in ECs during endothelialization, including anchoring, attachment, spreading, and motility. Notably, we found that higher RI values in attached ECs are associated with increased cell motility, suggesting that RI is a powerful indicator of cellular behavior that can be leveraged in endothelialization experiments.

Our results indicate that RI is a powerful indicator of the early stages of endothelialization that could be used in endothelialization experiments to gauge the affinity of cells to a matrix material, biomaterial, or medical device surface coating in a label-free and high-throughput manner. RI values of cells decreased during the initial stages of endothelialization, specifically during cell anchoring, attachment, and spreading. This decrease was followed by stabilization of RI values as cells established firm adhesions and formed monolayers. The precipitous drop in RI values may be attributed to changes in cell density and volume as cells spread out, as well as redistribution of cytoplasmic components. The correlation between RI stabilization and morphological changes, such as increased cell area and changes in cell solidity, reinforces the link between physical cell properties and their optical characteristics measured by HTM. Recent work has shown that HTM imaging can help identify mutation states of cells ^15^ or differentiation status ^16^, and our report adds further evidence that HTM can gain quantitative insights into the functional states of living cells.

The positive correlation between higher RI values and increased cell motility during endothelialization suggests that specific cellular components or structural organizations contribute to both higher RI and enhanced migratory behavior, such as mitochondria or lipid droplets. Mitochondria are of particular interest in ECs, as it was recently reported that the donation of mitochondria from mesenchymal stromal cells to ECs through cell-cell tunneling nanotubes is critical for effective re-endothelialization of damaged tissue ^17^. Furthermore, mitochondrial fission and fusion have been implicated in determining endothelial health by regulating ROS levels ^18^. Since direct visualization of mitochondrial dynamics is difficult using traditional microscopy, HTM offers a promising alternative to study EC energy production, reactive oxygen species signaling, and subcellular component interactions.

HTM also enables clear visualization of lipid droplets, which have recently been identified as contributors to endothelial cell dysfunction by impinging on nitric oxide and VCAM levels. ^19,20^ These organelles could be more abundant or reorganized in motile cells to meet the energetic and biosynthetic demands of migration. Cytoskeletal dynamics may also play a role, as rearrangements of actin filaments and microtubules during migration could affect cellular density and, consequently, RI measurements. The increased protein density in regions of active cytoskeletal remodeling could contribute to higher RI values observed in motile cells. Our findings align with previous studies that link cellular motility with metabolic activity and cytoskeletal organization.^21,22^ For instance, cells with higher metabolic rates often exhibit increased motility and may display altered distributions of organelles involved in energy production and lipid metabolism.

Combining HTM with fluorescence imaging enhances the analytical power by providing both structural and molecular information.^23^ While HTM offers detailed structural visualization and quantitative RI measurements, it lacks specificity at the protein level. Immunofluorescence complements HTM by allowing the localization of specific proteins within the cellular context provided by HTM. Our use of VE-cad staining exemplifies this synergy, as high RI values at cell borders did not directly correlate with VE-cadherin localization. This suggests that HTM and immunofluorescence provide complementary, orthogonal approaches to visualizing cell-cell interactions. While HTM highlights structural features and variations in RI, fluorescence imaging confirms the presence and localization of specific proteins. This multimodal imaging approach enhances our understanding of complex cellular processes by integrating quantitative structural data with molecular specificity. It allows for the validation of label-free observations and provides a more comprehensive picture of cellular function.

The principles demonstrated in this study can be applied to other cell types and biological processes. HTM could be utilized to study stem cell differentiation, cancer cell invasion, or immune cell activation. Applying HTM to disease models affecting ECs, such as atherosclerosis or diabetes, could provide valuable insights into pathological mechanisms and potential therapeutic targets. Experiments utilizing pharmacological agents or genetic manipulation to alter mitochondrial function or lipid metabolism could elucidate their roles in endothelialization, for example. Time-lapse HTM imaging during these interventions would provide dynamic insights into organelle function and cellular responses. The RI-based measurements in this study can be generalized to other cell types and other biological applications. Multimodal imaging, such as combining fluorescence with HTM, can provide valuable information like more quantitative protein species abundance, subcellular protein localization to specific organelles, and improved annotation of subcellular components observed via HTM.

### Limitations of the Study

Despite the advantages of HTM, several limitations must be acknowledged. A significant technical challenge is the segmentation of individual endothelial cells in HTM images. The similarity in RI values between the cytoplasm and the surrounding culture media complicates automated segmentation. Current AI-driven segmentation models, such as Cellpose and StarDist, underperform on HTM data due to these subtle differences and the complexity of the images. Manual segmentation is time-consuming and limits the throughput of data analysis. Developing specialized algorithms or training AI models specifically on HTM data of ECs could address this limitation and enhance the applicability of HTM in large-scale studies.^25^

Biological variability is another consideration. ECs exhibit inherent heterogeneity in behavior and response to environmental cues. While our study provides valuable insights, the generalizability of the findings may be influenced by this variability. Further studies with larger sample sizes and different EC types could strengthen the conclusions.

## Supporting information

Cell_A

## Acknowledgements

The research reported in this publication was supported by the National Institute of General Medical Sciences of the National Institutes of Health under Award Number SC2GM140991. We also gratefully acknowledge funding support from Schmidt Futures (W.L.).

## Methods

### Cell Culture

HUVECs (ATCCC) were cultured in FluoroBrite DMEM (Gibco) supplemented with 10% FBS (Gibco LifeTechnologies), 5% penicillin, streptomycin, and fungizone (Cytiva Hyclone antibiotic/antimycotic solution), and further enhanced with 5ng/mL rh VEGF, 5ng/mL rh EGF, 5ng/mL rh FGF basic, 15ng/mL rh IGF, 10 mM L-glutamine, 0.75 units/mL heparin sulfate, and 1 g/mL ascorbic acid (ATCC, PCS-100-041). Cells were cultured on collagen-coated (Gibco, A1048301) T75 flasks (Southern Labware, SKU 708013) at 37°C and 5% CO_2_. Media was changed every 2 days, and cells were passed at 80-90% confluency.

### Immunocytochemistry

HUVECs were cultured in 35 mm wells with tissue-culture treated glass bottoms (Ibidi, USA). Cells were fixed with 4% (v/v, in PBS) paraformaldehyde (Invitrogen, Image-It FB-002) for 15 minutes at room temperature. The cells were permeabilized with 0.1% (v/v, in PBS) Triton-X100 (Invitrogen, HFH10) for 10 minutes and subsequently blocked with 5% BSA (w/v, in PBS) with 0.05% (v/v) Tween-20 (Thermo Scientific Chemicals, J63711-AK) for 30 minutes at room temperature. The primary antibody VE-cadherin (D87F2) XP® Rabbit mAb #2500 (CellSignaling, 2500) was diluted 1:100 in the blocking buffer and incubated with the cells for 60 minutes at room temperature. The secondary antibody anti-rabbit IgG (H+L), F(ab’)2 Alexa Fluor 647 (CellSignaling, 4414) was diluted 1:500 and incubated for 30 minutes at room temperature in the dark. Nuclei were stained with DAPI (Ibidi, Cat.No: 50011). The samples were stored at 4℃ until imaging.

### CX-F Holotomographic Image Acquisition

The 3D Cell Explorer-Fluo CX-F (Nanolive SA, Switzerland) microscope, equipped with a 60X objective, was used to capture holotomography images. A z-stack of 30 μm with 312.5 nm slice thickness and 90 μm x-y length was acquired. Refractive index (RI) profiling involved capturing 96 z-slices at 1.7-second intervals and 4° increments. Lateral (x-y) and axial (z) resolutions were 200 nm and 400 nm, respectively. Auto-calibration accounted for the immersion medium (DMEM, RI: 1.3370). Fixed HUVECs were analyzed using DAPI and CY-5 filter cubes, with gain, exposure, and intensity parameters set according to the manufacturer’s recommendations (Table 1).

**TABLE 1:** CX-F Excitation Signal Parameters for Immunofluorescence Imaging.

| Channel | Exposure | Gain | Intensity |
| --- | --- | --- | --- |
| DAPI | 150 - 300 ms \| aimed 200 - 240 ms | 5 - 15 \| aimed ~ 10 | 5 - 20 \| aimed ~ 10 |
| Cy5 | 150 - 300 ms | 10 - 25 \| aimed below<br>20 | Preferred 10 - 25 \|<br>Aimed above 25 |

### CX-A Holotomographic Image Acquisition

Large area live imaging utilized the 3D Cell Explorer CX-A 96 Focus (Nanolive, UK). The CX-A’s optical specifications matched the CX-F, with the addition of a configurable field of view (FOV). Image stitching created larger 10x10 FOVs from standard 90x90 µm stacks. Atmospheric conditions were maintained using the CX-A’s incubation system (Tokai Hit, STX). A 10x10 grid scan was performed starting 4 hours post-seeding and images were acquired every 12.5 minutes for 16 hours.

### Image Analysis

Manual cell segmentation was performed using ImageJ FIJI to define cell outlines. The resulting ROIs were saved and used to measure the mean RI value and cell shape descriptors. Solidity, one such shape descriptor, is defined as the ratio between the area of a shape and the smallest possible bounding rectangle. The center of mass from these shapes were used for cell tracking analysis.

### Statistical Analysis

Spearman correlations were performed using the SciPy library in Python. Colocalization between fluorescence and RI channels was determined using the Costes colocalization method, using the ImageJ FIJI Coloc2 plugin.^24^ Time series data was smoothed using a 2-hour rolling window to calculate the mean and 95% confidence intervals.

## Notes

### Competing Interest Statement

The authors have declared no competing interest.

### Summary of Updates

Improved description of HTM as well as Figure descriptions.

## References

1. Kim, G. et al. Holotomography. Nature Reviews Methods Primers 4, 1–22 (2024).

2. Ghosh, B. & Agarwal, K. Viewing life without labels under optical microscopes. Commun Biol 6, 559 (2023).

3. Medina-Ramirez, I. E., Macias-Diaz, J. E., Masuoka-Ito, D. & Zapien, J. A. Holotomography and atomic force microscopy: a powerful combination to enhance cancer, microbiology and nanotoxicology research. Discov Nano 19, 64 (2024).

4. Fritsche, S., Fronek, F., Mach, R. L. & Steiger, M. G. Applicability of non-invasive and live-cell holotomographic imaging on fungi. J. Microbiol. Methods 224, 106983 (2024).

5. Suh, J. et al. Mitochondrial fragmentation and donut formation enhance mitochondrial secretion to promote osteogenesis. Cell Metab. 35, 345–360.e7 (2023).

6. Kim, H., Oh, S., Lee, S., Lee, K. S. & Park, Y. Recent advances in label-free imaging and quantification techniques for the study of lipid droplets in cells. Curr. Opin. Cell Biol. 87, 102342 (2024).

7. Sandoz, P. A., Tremblay, C., van der Goot, F. G. & Frechin, M. Image-based analysis of living mammalian cells using label-free 3D refractive index maps reveals new organelle dynamics and dry mass flux. PLoS Biol. 17, e3000553 (2019).

8. Hayakawa, E. H., Yamaguchi, K., Mori, M. & Nardone, G. Real-time cholesterol sorting in Plasmodium falciparum-erythrocytes as revealed by 3D label-free imaging. Sci. Rep. 10, 2794 (2020).

9. Huang, H. et al. Quantitative label-free digital holographic imaging of cardiomyocyte optical volume, nucleation, and cell division. J. Mol. Cell. Cardiol. 196, 94–104 (2024).

10. Jana, S. Endothelialization of cardiovascular devices. Acta Biomater. 99, 53–71 (2019).

11. Wolfe, J. T., Shradhanjali, A. & Tefft, B. J. Strategies for Improving Endothelial Cell Adhesion to Blood-Contacting Medical Devices. Tissue Eng. Part B Rev. 28, 1067–1092 (2022).

12. Zhao, J. & Feng, Y. Surface Engineering of Cardiovascular Devices for Improved Hemocompatibility and Rapid Endothelialization. Adv. Healthc. Mater. 9, e2000920 (2020).

13. Jerka, D. et al. Unraveling Endothelial Cell Migration: Insights into Fundamental Forces, Inflammation, Biomaterial Applications, and Tissue Regeneration Strategies. ACS Appl Bio Mater 7, 2054–2069 (2024).

14. Reglero-Real, N., Colom, B., Bodkin, J. V. & Nourshargh, S. Endothelial cell junctional adhesion molecules: Role and regulation of expression in inflammation. Arterioscler. Thromb. Vasc. Biol. 36, 2048–2057 (2016).

15. Kim, H. et al. Integrating holotomography and deep learning for rapid detection of NPM1 mutations in AML. Sci. Rep. 14, 23780 (2024).

16. Sbrana, F. et al. Label-free three-dimensional imaging and quantitative analysis of living fibroblasts and myofibroblasts by holotomographic microscopy. Microsc. Res. Tech. 87, 2757–2773 (2024).

17. Lin, R.-Z. et al. Mitochondrial transfer mediates endothelial cell engraftment through mitophagy. Nature (2024) doi:10.1038/s41586-024-07340-0.

18. Kluge, M. A., Fetterman, J. L. & Vita, J. A. Mitochondria and endothelial function. Circ. Res. 112, 1171–1188 (2013).

19. Jaffe, I. Z. & Karumanchi, S. A. Lipid droplets in the endothelium: The missing link between metabolic syndrome and cardiovascular disease? J. Clin. Invest. 134, (2024).

20. Wazir, M. et al. Lipid disorders and cardiovascular risk: A comprehensive analysis of current perspectives. Cureus 15, e51395 (2023).

21. Cunniff, B., McKenzie, A. J., Heintz, N. H. & Howe, A. K. AMPK activity regulates trafficking of mitochondria to the leading edge during cell migration and matrix invasion. Mol. Biol. Cell 27, 2662–2674 (2016).

22. Shiraishi, T. et al. Glycolysis is the primary bioenergetic pathway for cell motility and cytoskeletal remodeling in human prostate and breast cancer cells. Oncotarget 6, 130–143 (2015).

23. Larrazabal, C., Hermosilla, C., Taubert, A. & Conejeros, I. 3D holotomographic monitoring of Ca++ dynamics during ionophore-induced Neospora caninum tachyzoite egress from primary bovine host endothelial cells. Parasitol. Res. 121, 1169–1177 (2022).

24. Costes, S. V. et al. Automatic and quantitative measurement of protein-protein colocalization in live cells. Biophys. J. 86, 3993–4003 (2004).

25. Michael, R., Modirzadeh, T., Issa, T. B. & Jurney, P. Label-Free Visualization and Segmentation of Endothelial Cell Mitochondria Using Holotomographic Microscopy and U-Net. 2024.11.26.625487 Preprint at 10.1101/2024.11.26.625487 (2024).

