## Supplementary figures and images for "Holotomographic microscopy reveals label-free quantitative dynamics of endothelial cells during endothelialization"

### Cell_A

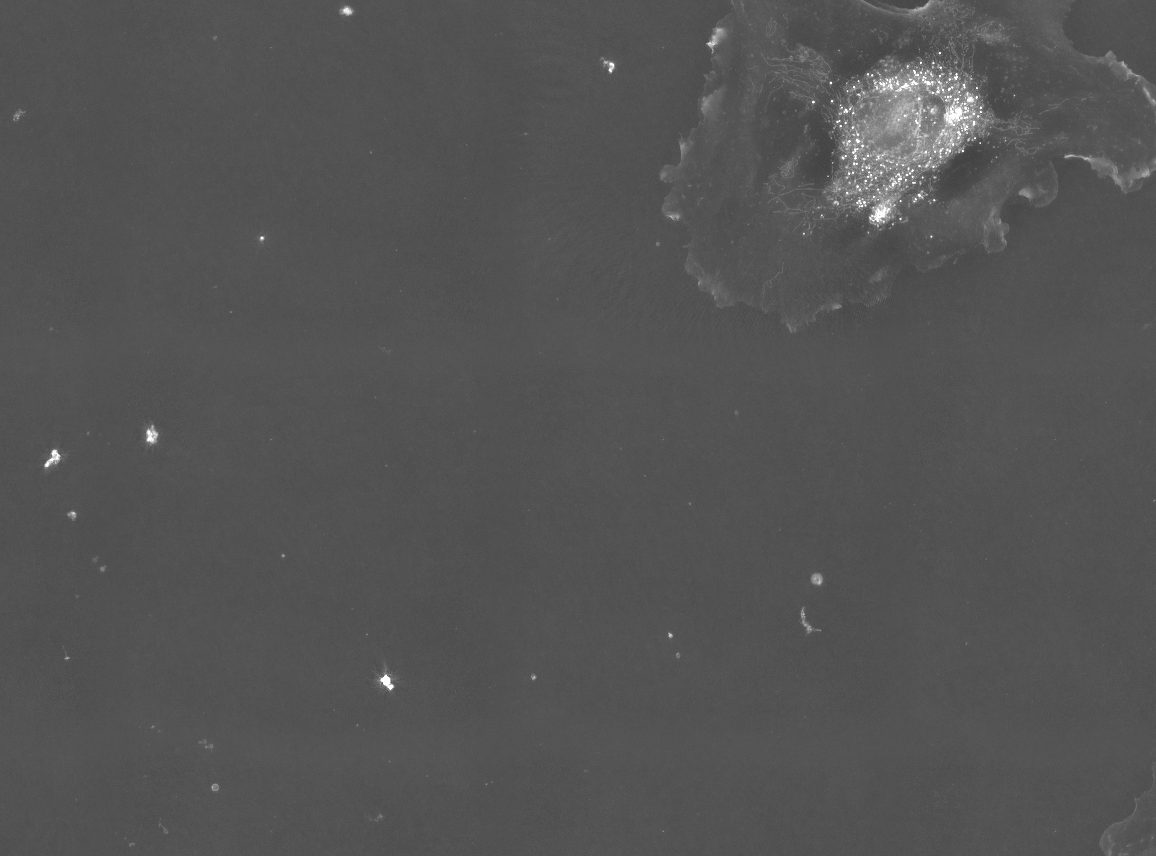
